# Acclimation to thermal variability increases the intensity of activity and alters the activity window in the temperate dung beetle *Onthophagus taurus*

**DOI:** 10.64898/2026.03.03.708744

**Authors:** Alexander J Coverley, Kimberly S. Sheldon, Katie E. Marshall

## Abstract

1. Ectotherms in thermally variable environments mediate energy expenditure through both physiological and behavioural responses. However, many studies focus on constant temperature acclimation, and few consider behaviour and physiology in unison. It is unclear how acclimation to thermal variability affects locomotory choices, activity timing, and performance across daily thermal cycles.
2. We investigated the effects of thermal variability in the temperate dung beetle *Onthophagus taurus*. Following acclimation to a low amplitude (22°C ± 2°C) or a high amplitude (22°C ± 10°C) temperature regime, we measured behaviour and metabolic rate across temperatures. We hypothesised that *O. taurus* adjusts its locomotive strategy and search window when kept in high amplitude fluctuating temperatures to reduce energy loss associated with high temperature exposure.
3. We found that differences in energy expenditure were determined by propensity for flight which differed between acclimation treatments, particularly at intermediate temperatures. We also found that, following acclimation to a high amplitude of thermal variability, *O. taurus* exhibited a greater intensity of activity over a narrower window of time, and *O. taurus* acclimated to a low amplitude of thermal variability showed nocturnal activity.
4. We then used the data to model activity through the growing season over five years. Biophysical models were built using NicheMapR Microclimate and Ectotherm functions to test the length of potential searching time across seasons, the temperatures individuals are exposed, and locomotive strategy. Model outputs showed that acclimation to higher amplitudes of thermal variability increased accumulated degree-hours of activity relative to the low variability acclimation group. Individuals acclimated to higher amplitudes of thermal variability showed greater accumulated degree-hours in spring and fall, but exhibited shorter periods of activity during summer, with the model predicting increased opportunities for flight. Comparatively, *O. taurus* from the low variability acclimation treatment showed increased night activity in summer but did not fly.

## Introduction

Most insect species are ectothermic, meaning their body temperature is largely driven by external conditions (Heinrich, 2013). As such, temperature restricts foraging and mate-seeking opportunities in mobile species (Leith et al., 2021; Willmer 1983). However, morphological and physiological traits such as body size, colouration, and the amount of metabolic heat production due to muscle contraction can allow an insect’s body temperature to differ from the surrounding air (Clusella-Trullas & Nielsen, 2020; Heinrich, 1974; Harris et al., 2012). In temperate environments, air temperature fluctuates over daily and seasonal time scales (Dillon et al., 2016), limiting activity and performance.

Every species has a suitable thermal range, described here as the thermal activity window, that defines the period in which searching behaviours such as locating resources and mates are possible (Everatt et al., 2013). Within a daily thermal cycle, environmental temperatures outside of the suitable thermal range can occur because of both high and low temperatures. Thus, high amplitude fluctuations in temperature can constrain behaviour at multiple parts of the daily thermal cycle. In temperate diurnal species, activity is generally possible for longer periods during the summer relative to winter due to warmer temperatures (Wolda, 1988). However, midday high summer temperatures can exceed the upper thermal limits of the organism, in some cases leading to bimodal activity windows (Xu et al., 2021; Lobo & Cuesta 2021). To avoid undesirable temperatures, motile insects use behavioural thermoregulatory responses such as retreat.

However, this can reduce search time and therefore energy acquisition (Carter et al., 2025). Across the range of temperatures where locomotion is possible, shifts in temperature mediate performance, altering movement speed and associated energy demand.

Thermal performance curves (TPCs) describe the effect of temperature on biological rates such as locomotion and metabolism (Sinclair et al., 2016). These rates canonically rise in an accelerating fashion towards an inflection point, decelerate towards the thermal optimum, then decline rapidly at temperatures exceeding the optimum as biological processes begin to break down. Jensen’s inequality is a property of nonlinear functions that, when applied to TPCs, means that the mean performance for a biological rate differs from the performance at the mean temperature (Ruel et al., 1999; Denny, 2017). The amplitude of fluctuation across a daily thermal cycle affects energy expenditure independently of mean temperature as the rate of increase in metabolic rate is non-linear. Consequently, applying mean temperature as a proxy for mean metabolic rate can lead to underpredicting energy expenditure in thermally variable environments.

In temperate environments where temperature can fluctuate rapidly, timing locomotory activity to peak at suboptimal temperatures can increase fitness by reducing the risk of accidentally straying into temperatures in the area of rapid decline in performance (Dowd et al., 2015; Martin & Huey, 2008), although strategy can vary across species. For example, in Mediterranean ant communities, activity periods differ between heat-intolerant species that forage at lower temperatures and heat-tolerant species that forage nearer their critical thermal maximum with greater performance and mortality risk (Cerdá et al., 1998). Within a daily thermal cycle, trade-offs exist based on when searching occurs across high and low temperatures with high temperature searching activity being associated with greater energetic demand and increased performance, but greater interactions with resources while foraging (Colinet et al., 2015; Hannigan et al., 2023).

Of all the forms of locomotion for foraging, flight is the most energetically expensive and requires higher body temperatures than walking. To fly, insects must first complete a pre-flight warm-up by contracting thoracic muscles to raise thoracic temperature, and consequently, metabolic rate (Casey et al., 1981). In the mosquito *Aedes aegypti*, crawling ceases at 10 °C, and while flight can occur just above this temperature, it cannot be sustained until a higher threshold of 15 °C is met (Rowley & Graham, 1968). Similarly, Verdú et al., (2006) found thermal constraints on the minimum and maximum flight temperature across multiple taxa of dung beetles, with a minimum body temperature of 25 °C necessary to achieve flight. Maintaining endogenous heat production is energetically expensive, and metabolic rate during flight can be between 50 and 100 times higher than resting metabolic rate (Kammer & Heinrich, 1978). Therefore, the shape of the TPC for flight metabolic rate differs greatly from that of crawling because of the high energetic cost across a narrower range of temperatures (Glass & Harrison 2022), reducing opportunities for flight across daily and seasonal timescales.

The energy expended on locomoting during foraging is therefore dependent on the length of time the organism forages, the temperature while foraging, and the locomotive method used. If the activity window of an individual shifts, then the daily heat and energy balance may also change. For facultative fliers (animals that can effectively locomote via either flight or other means), low temperature can constrain flight while crawling remains a viable alternative, reducing search rate and dispersal (Boiteau, 2000; Caprio & Grafiusm 1990; & Wegorek, 1963). As these locomotory behaviours modulate responses to temperature as a consequence of previous exposure, a biophysical modelling approach is well-suited to predicting these differences in activity timing, length, and accumulated degree hours. A biophysical model predicts thermoregulatory responses at the organismal level by integrating the behavioural and physiological traits of an organism with the thermal conditions of its environment by calculating heating exchange between the two (Porter & Gates, 1969; Porter et al., 1973). The available surface temperatures for activity are dependent on behavioural parameters set so that physiological constraints – including thermodynamic constraints – limit when activity is possible. For example, when the organism retreats underground to avoid unfavourable temperatures, it is unable to forage, shortening its activity window, but giving a realistic representation of total available searching time within its environment and the organism’s energy budget (Angilleta, 2001).

All of the above-described thermal limits and relationships are plastic, meaning they can be shifted in response to changing environmental conditions. Some insect species exhibit seasonal plasticity in the lower temperature threshold for flight. For example, the mountain pine beetle adjusts its minimum flight temperature between spring (18.6 °C) and autumn (below 13 °C) (Gaylord et al., 2008). Acclimation to the breadth of experienced temperatures in a daily cycle can also alter performance, fecundity, and metabolic rate. For example, acclimation to thermal variability increases flight performance, cold tolerance, and egg laying capacity in the false codling moth *Thaumatotibia leucotreta* (Huisamen et al., 2022). Pupae of the dung beetle *Onthophagus taurus* possess no effective locomotive capacity, and acclimation to thermal variability reduces the thermal sensitivity of metabolic rate (Fleming et al., 2021). However, despite many studies examining the effect of thermal variation on metabolic rate and thermal limits, it remains unclear whether acclimation to thermal variation influences locomotory choices, which could potentially feedback on the ability to behaviourally thermoregulate and resulting energetic costs.

This study used the widely studied dung beetle *Onthophagus taurus* (Casasa et al., 2017; Emlen & Nijhout, 2001; Hunt et al., 2002; Moczek & Emlen, 1999; Nijhout, 2003; Fleming et al., 2024) to test whether acclimation to thermal variability affects locomotive choices and their associated energetic costs. We first hypothesized that *O. taurus* adjusts its locomotive strategy and search window when kept in high amplitude fluctuating temperatures to reduce energy loss associated with high temperature exposure. If this is true, we predict that t 1) under increased amplitudes of thermal variability, daily activity will be concentrated into a shorter period of time and time spent above ground will be reduced; and that acclimation to thermal variability 2) increases the time spent locomoting and 3) suppresses metabolic rate. We found that *O. taurus* acclimated to a high amplitude of thermal variability have a greater intensity of activity over a narrower window of time. However, *O. taurus* did not exhibit metabolic suppression, rather differences in metabolic rate between treatments was determined by differences in flight behaviour, most prominent at intermediate temperatures. We also found that beetles acclimated to a low amplitude of thermal variability showed nocturnal activity but reduced overall activity. We incorporated these results into a biophysical model to predict how acclimation to thermal variability affects activity window, locomotive opportunities and accumulated searching degree hours in the wild. Of the predictions that daily accumulated degree-hours and activity window would be reduced in *O. taurus* acclimated to thermal variability, only the latter held true. Accumulated degree hours were higher in those acclimated to variability and substantially higher than the constant temperature acclimated beetles during spring and fall.

## Methods

### Onthophagus taurus

*Onthophagus taurus* (Coleopetera: Scarabaeidae) is a small (<150 mg), generalist dung eating beetle and a Mediterranean species that has been introduced to Australia, Hawaii, and the eastern United States. It is a facultative flyer that burrows to store dung, bury brood balls, and thermoregulate, and surfaces to conduct searching behaviour for foraging and mating (Halffter & Edmonds, 1982; Snell-Rood et al., 2016). Activity extends through spring and autumn, sometimes with two generations a year, and late-summer progeny overwinter in the adult stage (Floate et al., 2017). At high temperatures, *O. taurus* will commonly remain at dung pats feeding for over a day before flying to locate fresh dung (Hunt et al., 1999). It also exhibits plasticity in thermal physiology, showing higher heat acclimation ability than a congener (Mamantov & Sheldon, 2025) as well as metabolic plasticity, reducing its resting metabolic rate under greater amplitudes of thermal variability (Carter & Sheldon, 2020).

### Collection and acclimation

We collected Female *Onthophagus taurus* from cowpats in fields around Knoxville, Tennessee (USA) in May and June 2023. We acclimated forty-three *O. taurus* to a low variation temperature regime of 22°C ± 2°C for four days, with a 16:8 light dark cycle (hereafter L:D cycle), in an incubator (PHCbi: MIR-554-PA, Japan). Following this, we assigned beetles to one of two temperature fluctuation treatments: a high fluctuation treatment of 22°C ± 10°C (n=23) and a low fluctuation treatment of 22°C ± 2°C (n=20), all with a 16:8 L:D cycle and using the same incubator model.

We kept beetles in 237 mL containers with 120g of a compressed mixture of three parts autoclaved soil to two parts autoclaved sand and fed autoclaved cow dung once every four days and again on the day of the behavioural assay. The dung was collected from cow pastures belonging to Cruze Farm Dairy, Knoxville. We weighed all beetles at the start and end of the treatment period. After six days of fluctuation treatments, we transferred beetles into a fresh container and returned to the same incubator. Three beetles perished during the fluctuation treatment period: one from the high fluctuation treatment and two from the low fluctuation treatment.

**TABLE 1.** Replication statement. For the scale of inferences made, the scale at which treatment was applied, and the number of replicates for each level of treatment.

| Scale of inference | Scale at which treatment was applied | Number of replicates |
| --- | --- | --- |
| Individuals | Females | $22^{\circ}\text{C} \pm 2^{\circ}\text{C}$: 14<br>$22^{\circ}\text{C} \pm 10^{\circ}\text{C}$: 15 |

### 24-hour behavioural assay

To record activity and behaviour, we monitored the beetles for 24 hours in their fluctuation treatments using a 1080P digital video camera (Infra-Red, 2.7K, 30 MP, 2688 x 1520 pixels, 16x zoom) inside the incubator. We used the camera’s zoom function to maximise the size of the beetles in the footage, and we used night mode to allow for recordings during the dark. We placed a matte brown cardboard background behind the containers to reduce reflection. One individual was partially obscured during the recording and removed from further consideration, resulting in 14 individuals being used for the low fluctuation treatment group and 15 for the high fluctuation treatment group.

We reviewed the full videos, advancing in 10 second increments until the beetle emerged onto the soil surface, and recorded the duration of time the beetle was still and active. Movement was defined as any period the beetle was in motion, including emerging, walking, digging, or climbing. We only scored behaviours performed on the soil surface. Time was binned into two-hour intervals that corresponded to periods of stable incubator temperature (Table S1).

### Metabolic rate measurements

We made metabolic rate measurements using a flowthrough respirometry system with a Li-7000 carbon dioxide gas analyser (LI-COR BIOSCIENCES, Lincoln, Nebraska, USA). Here, measurements followed seven days of acclimation to the experimental temperature regime (including the day of the behavioural assay). We weighed beetles to the nearest mg and individually placed them in a 500 mL respirometry chamber, inside a water bath (Neslab RTE7, Newington, New Hampshire, USA) to control temperature. We placed an identical empty chamber next to it with a Type T thermocouple at the bottom and top of the chamber, connected to a Picolog interface (Pico Technology, Cambridgeshire, UK) to record air temperature using Picolog software (Pico Technology, Cambridgeshire, UK). Dry Ultrazero air (Airgas USA, LLC; 20-22% O_2_, 2%H2, 0.1% THC, 1% CO_2_, 1% CO, 74.9-76.9% N_2_) was pushed into the chamber at 400 mL/min, with flow rate controlled by a Mass Flow Controller (ALICAT Scientific, Tuscon, Arizona, USA). Baseline readings were taken over five minutes before placing the beetle in the chamber. Beetles were left in the chamber while gas flowed through the system without light for a period of 22 minutes at 22°C to ensure the removal of any excess CO _2_ from opening the chamber. At the beginning of the measurement, a 2-cell incandescent flashlight with a 99 metre beam distance was held over the chamber to simulate sunlight and make the beetle easier to see. The water bath containing the chamber was held at 22°C for 10 minutes. The bath temperature was then ramped to 25°C over a period of 8 minutes and held for 10 minutes, then ramped over 8 minutes to 30°C and held for 10 minutes. Behaviour and carbon dioxide production rate were recorded for each ten minutes at a constant temperature.

Activity was recorded continuously during the assay as one of the following options: still, walking, and flying. If the beetle was on its back, having fallen during flight, we noted the amount of time it was upside down and this period was excluded from the analysis.

To measure metabolic rate, we first converted CO _2_ measurements from parts per million concentrations to mL/min of CO _2_ using Equation (1):

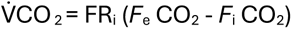

**Equation (1)**: where V̇CO _2_ is the rate of CO _2_ production in mL/min, *F*_i_ CO_2_ is the incurrent fractional CO_2_ concentration, *F*_e_ CO_2_ is the excurrent fractional CO_2_ concentration, FR_i_ is the incurrent flow rate in mL/min. Output was matched to both the corresponding temperatures recorded by the Picolog software and the observed behaviour.

### Statistical analysis

We first used a linear model (“lm” function in base R; R version 4.2.2) to confirm there was no significant difference in beetle masses at the beginning of the acclimation period using initial mass as a response variable and acclimation treatment as a predictor. A second linear model was used to determine how acclimation to thermal variability affects body mass for beetles acclimated to 7 days at near-constant or variable treatments. The explanatory variables were the mass at the beginning of the fluctuation treatment period, the fluctuation treatment, and their interaction.

For the 24-hour behavioural assays, we compared either time spent moving or time spent still above ground between the experimental groups. Specifically, we used two different Generalised Linear Mixed Model (GLMM) with loglink error distributions using the glmmTMB package (Brooks et al., 2017). This was appropriate for a high number of zeroes, which arose when a behaviour did not occur within the two-hour window during which temperature was held constant, and allowed us to fit a zero inflation probability term “Zi =∼1” to account for covariance between light and temperature cycles in the model (Brooks et al., 2017). For both models – time spent still above ground and time spent moving above ground–the response variables were fluctuation treatment, incubator temperature, L:D cycle, and the interaction between temperature and L:D cycle. Beetle ID was treated as a random factor to account for non-independence of data points. Each model was then compared with a simpler model that excluded fluctuation treatment, a null model, and a model with a Poisson distribution using Aikake’s information criterion (AIC) comparisons (Table S2 & Table S3).

For behaviour during respirometry, we analysed the effect of fluctuation treatment on time spent on each locomotive behaviour across the ramped water temperature (hereafter, trial temperature) in three separate models to decrease model complexity (for still, crawling, and flying). Data points of when beetles were upside down after hitting the glass while flying were omitted from the study. For each locomotive behaviour, we again used a GLMM fitted using the glmmTMB package (Brooks et al., 2017) in R version 4.2.2. The proportion of time spent on each form of locomotion was modelled with a negative binomial distribution (nbinom2) to account for overdispersion. This is because pre- flight warm up causes clustering of stillness and flight events, and thus locomotive type is not independent of previous events. The models for time spent in still and crawling behaviours included fluctuation treatment, trial temperature, and mass as fixed effects and beetle ID as a random effect because of repeated measures across temperature. The model for flight was adapted in two ways.

First, as the minimum threshold for flight was higher than the lowest assayed trial temperature, a zero inflating term (Zi = ∼1) was included, which incorporates a submodel for the probability of a value other than zero being impossible under some conditions (Brooks et al., 2017). Secondly, the fluctuation treatment and trial temperature were treated as interacting terms, as we expected flight occurrences to increase significantly under higher temperatures. AIC scores were compared with a simpler model excluding the fluctuation treatment and the model with the lowest AIC score was selected (Flight, Table S4; Walking, Table S5; and Still, Table S6).

To test the hypothesis that beetles acclimated to higher amplitudes of thermal variability would reduce energy expenditure, CO _2_ production was first compared between fluctuation treatment groups across trial temperature. Linear mixed-effects models were fitted using the lme4 package (Bates et al., 2015). Total CO _2_ production – across all behaviours – at a given trial temperature was used as the response variable, to capture overall energy expenditure and to account for increased energy expenditure during pre-flight warm up and post-flight. The explanatory variables were body mass at the start of the respirometry assay, time spent flying, the interaction between time spent flying and fluctuation treatment, and the trial temperature. Beetle ID was included as a random effect. Model selection was performed by comparing AIC scores with simpler models excluding terms (Table S6).

Metabolic rate was compared for each behaviour and trial temperature combination between experimental groups. Factors affecting V̇CO _2_ in flying beetles were tested by fitting a linear mixed- effects model with individual ID as a random effect (Bates et al., 2015). As there were few repeated measures of the same individuals across trial temperature, no evidence suggesting individual- specific effects, and poor dispersion, ID was removed to improve model fit, before fitting a linear model and ANOVAs (base R; R version 4.2.2). The fixed effects were fluctuation treatment, trial temperature, mass at the start of the trial, and the count of flying behaviour. The count of flying behaviour was included to test whether the time spent in flight affects flight metabolic rate. Model selection was performed by comparing AIC scores with simpler models excluding terms (Table S7).

To test factors affecting V̇CO _2_ in beetles while they were walking, a GLMM was fitted with tweedie type distribution and log link (Brooks et al., 2017). The tweedie type distribution was more appropriate as there was greater increase in time spent crawling with trial temperature. The response variables were fluctuation treatment, trial temperature, and mass at the start of the trial. Model selection was performed by comparing AIC scores with simpler models excluding terms (Table S8).

To test factors affecting V̇CO _2_ in beetles while they remained still, a GLMM was fitted with a gamma type distribution and log link (Brooks et al., 2017). This distribution was selected based the right-skewed distribution. Model selection was performed by comparing AIC scores with simpler models excluding terms (Table S9). The response variables were fluctuation treatment, trial temperature, and mass at the start of the trial.

### Modelling

The R package NicheMapR incorporates a microclimate model that estimates temperatures above ground and below the soil surface using heat and mass balance equations in conjunction with local abiotic and biotic components such as altitude, windspeed, and soil properties. The outputs of the microclime model are used in the NicheMapR ectotherm model to estimate the body temperature of the organism incorporating behavioural thermoregulatory strategies such as burrowing, diel activity patterns, and the distance between the organism and the soil surface (Kearney & Porter 2017; Kearney & Porter 2020).

In order to test whether acclimation to thermal variability affected energy expenditure in *O. taurus,* daily accumulated degree hours and activity window were compared in a biophysical model. We predicted reduced hours available for searching and lower accumulated degree hours in *O. taurus* acclimated to high amplitude fluctuating temperatures compared with low amplitude fluctuating temperatures when modelling behaviour under field conditions. Winter days from DOY: 0-90 and 300-360 were excluded from the analysis as a conservative estimate of when the animal would be overwintering based on field data on emergence timings (Wassmer, 2020) and the minimum thermal threshold for development being 15.5°C (Floate et al., 2014). The model outputs were used to estimate the daily amount of time that searching is possible, conditions are met to search, and flight is possible, and the accumulated daily degree hours of activity.

The microclimate model was built in NicheMapR using the Micro_NCEP function (Kearney & Porter, 2017) at an hourly resolution (see supplemental section: *details of biophysical models* and Table S10 for model structure). Recent years with available hourly data were selected and separate yearly simulations were run from 2018 to 2023. The selected site was Knoxville Airport, Tennessee, US (35.81801°, -83.98573°) as the closest site to where the beetles were collected and where hourly weather data is available for the selected years. Weather data was obtained from the NOAA’s Global Historical Climatology Network-Hourly database (Menne 2023).

The Ectotherm function in NicheMapR was run for each individual beetle in each yearly simulation. Individual level data from the 24-hour behavioural recordings incorporated into the model included wet weight, whether the beetle was active at night, and maximum foraging temperature. Maximum foraging temperature was determined by the maximum temperature at which each individual was active during the behavioural trials. Although the beetles were exposed to different fluctuation treatments during the 24 hours, the high fluctuation group had no individuals active at the maximum temperature, and the low fluctuation group had only one individual active at the maximum temperature. Therefore, the assumption was made that this was the maximum temperature at which activity would occur. As individuals in the low fluctuation group were active at the lowest assayed temperature of 20 °C, we set the minimum foraging temperature as the lower consistent temperature at which activity was recorded of 18 °C found in the high fluctuation group. This temperature is 2.5 °C above the minimum temperature required for the organism to develop into an adult found in the literature (Floate et al., 2015). Other forcing variables relating to *O. taurus* physiology necessary to calculate body temperature were taken from the literature (see supplemental section: *details of biophysical models* and Table S11 for model variables and limitations).

### Statistics

Model outputs were combined with the individual minimum flight temperature recorded during respirometry as an estimate of the lowest temperature at which the beetle would fly in the field. These were used to quantify the amount of the daily foraging time the individuals could fly over the course of the study period to calculate the degree hours available for foraging using equation 2:

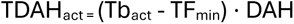

Equation 2: where TDAH_act_ is the total degree hours of activity per day. Tb_act_ is the hourly estimates of body temperature while active. TF_min_ minimum foraging temperature as determined by the 24-hour behavioural trials. DAH is the daily activity hours.

To test differences in accumulated degree hours during activity between fluctuation treatments, we fitted a GLMM with tweedie type distribution and loglink (Brooks et al., 2017). Daily degree hours spent searching was modelled with the response variables day of year, fluctuation treatment, the interaction between the day and fluctuation treatment, with the random effects of year and beetle ID. The term day of year included a second-order polynomial term to account for seasonal shifts in temperature. Model selection was performed by comparing AIC scores with simpler models excluding terms (Table S12).

## Results

### Body mass

The surviving 36 female *O. taurus* were weighed after a 4-day acclimation to 22°C ± 2°C, then reweighed after a 7-day temperature fluctuation treatment of either low (22°C ± 2°C) or high (22°C ± 10°C) fluctuation. *Onthophagus taurus* in the high fluctuation treatment had a slightly higher average initial mass (84 ± 4 mg) compared to those in the low fluctuation treatment (79 ± 4 mg), but this difference was not statistically significant (*F*_1,30_ = 0.720, *p* = 0.403). Initial mass was a strong predictor of mass at the end of the 7-day fluctuation treatment (*F*_1,28_ = 56.238, *p* < 0.001), while fluctuation treatment (*F*_1,28_ = 1.658, *p* = 0.208), and the interaction between fluctuation treatment and initial mass (*F*_1,28_ = 0.885, *p* = 0.335) were not significant predictors.

### Behaviour over 24 hours

Time spent crawling increased as incubator temperatures rose (χ^2^ _(1)_ =14.147, P < 0.005) with only one individual crawling during the part of the thermal cycle where temperatures declined. The LD cycle had a strong effect on time spent crawling with high levels of activity occurring under light exposure (χ^2^ _(1)_ =5.940, *P* = 0.015). The interaction between incubator temperature and the LD cycle was also a significant term where nighttime and daytime activity were both more common as temperatures rose (χ^2^ _(1)_ =6.323, *P* = 0.012). The beetles in the high fluctuation treatment spent a significantly higher proportion of time crawling when accounting for temperature, ID, and LD cycle (χ^2^ _(1)_ =3.889, *P* = 0.049). Despite this, the high fluctuation group only showed one instance of nocturnal activity and were constrained to a more limited activity time. The incubator temperatures in which the high fluctuation group were active exceeded the temperatures to which the low fluctuation group were exposed. However, the decline in activity at the lower and upper thermal range in the low fluctuation group suggests the low fluctuation group similarly reduces activity at lower temperatures during the thermal cycle. Four individuals in the high fluctuation group did not surface at all during the 24-hour period.

For time spent still while on the surface, the full model including acclimation treatment was not supported over the reduced model. Incubator temperature had the greatest explanatory power with stillness on the surface occurring more frequently as temperatures rose than when they declined (χ^2^ _(1)_ =5.901, *P* = 0.015). LD cycles explained some of the variation in occurrences with stillness on the surface being less common at night (χ^2^ _(1)_ =3.756, *P* = 0.053). The interaction between incubator temperature and LD cycles retained its significance because of the reasons addressed above (χ^2^ _(1)_ =3.756, *P* = 0.053). Only four beetles in the high fluctuation group were still while on the soil surface – and each of them only once – compared to twelve individuals in the low fluctuation group.

However, there was likely insufficient occurrence data for significant power.

**Figure 1:**
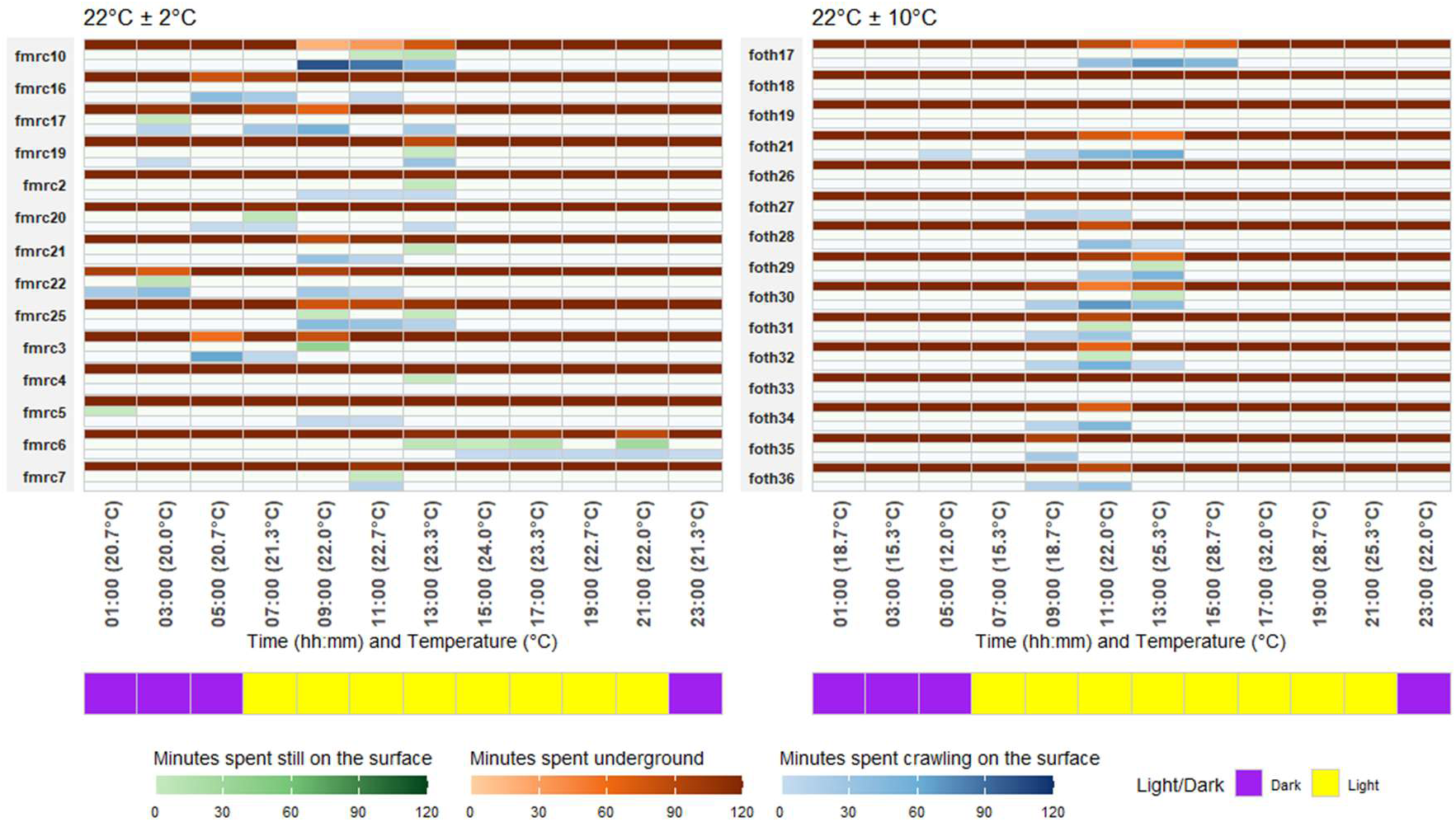
Activity budgets of female *Onthophagus taurus* over a 24-hour thermal cycle following a week of low (22°C ± 2°C) or high (22°C ± 10°C) fluctuation treatments. The X-axis shows time of day (2-hour bins) and associated incubator temperatures. The Y-axis shows the beetle ID. Colour scales show the number of minutes each behaviour occurred during each two-hour period. Green indicates the time spent still while on the surface, brown indicates time spent underground, and blue indicates time spent walking on the surface; if the behaviour did not occur at all in each two-hour period, then the box will be white. The purple and yellow bar shows the incubator light/dark cycle. Statistics presented in text.

### Behaviour during the respirometry trial

Fluctuation treatment affected occurrences of locomotive behaviour with high fluctuations being associated with greater activity (Figure 2). Fluctuation treatment had a significant effect on time spent still during respirometry measures, with those in the high fluctuation treatment spending less time sitting still across all tested temperatures (χ^2^ _(1)_ =7.532, *P* = 0.006). Trial temperature in the respirometry chamber was not a significant predictor of time spent still (χ^2^ _(1)_ =0.656, *P* = 0.798), and neither was mass (χ^2^ _(1)_ =0.110, *P* = 0.740). Time spent walking was solely explained by fluctuation treatment (χ^2^ _(1)_ =6.926, *P* = 0.008), but not trial temperature or mass (χ^2^ _(1)_ =0.517, *P* = 0.472; χ^2^ _(1)_ =0.225, *P* = 0.635). Trial temperature was a significant predictor of time spent flying (χ^2^ _(1)_ =14.868, *P* <0.005), with no instances of flight occurring below 25°C. For 32 beetles, the percent of individuals flying and time spent in flight increased from 19% of beetles at 25°C to 69% at 30°C. All individual beetles that flew at 25°C also flew at 30°C. Mass approached significance in explaining flight time with the seconds spent in flight increasing with mass when excluding those that did not fly at all (χ^2^_(1)_ =2.847, *P* = 0.092). Time spent flying was longer in those from the high fluctuation treatment at 25°C and 30°C (χ^2^ _(1)_ =6.063, *P* = 0.014). The interaction between trial temperature and fluctuation treatment was a predictor of time spent in flight (χ^2^ _(1)_ =5.478, *P* = 0.019). Differences in time spent flying between fluctuation treatments were more apparent at 25°C than 30°C (Figure 2). In summary, exposure to the higher fluctuation treatment increased the likelihood of flight at the lower thermal threshold for flight and reduced the amount of time spent still.

**Figure 2:**
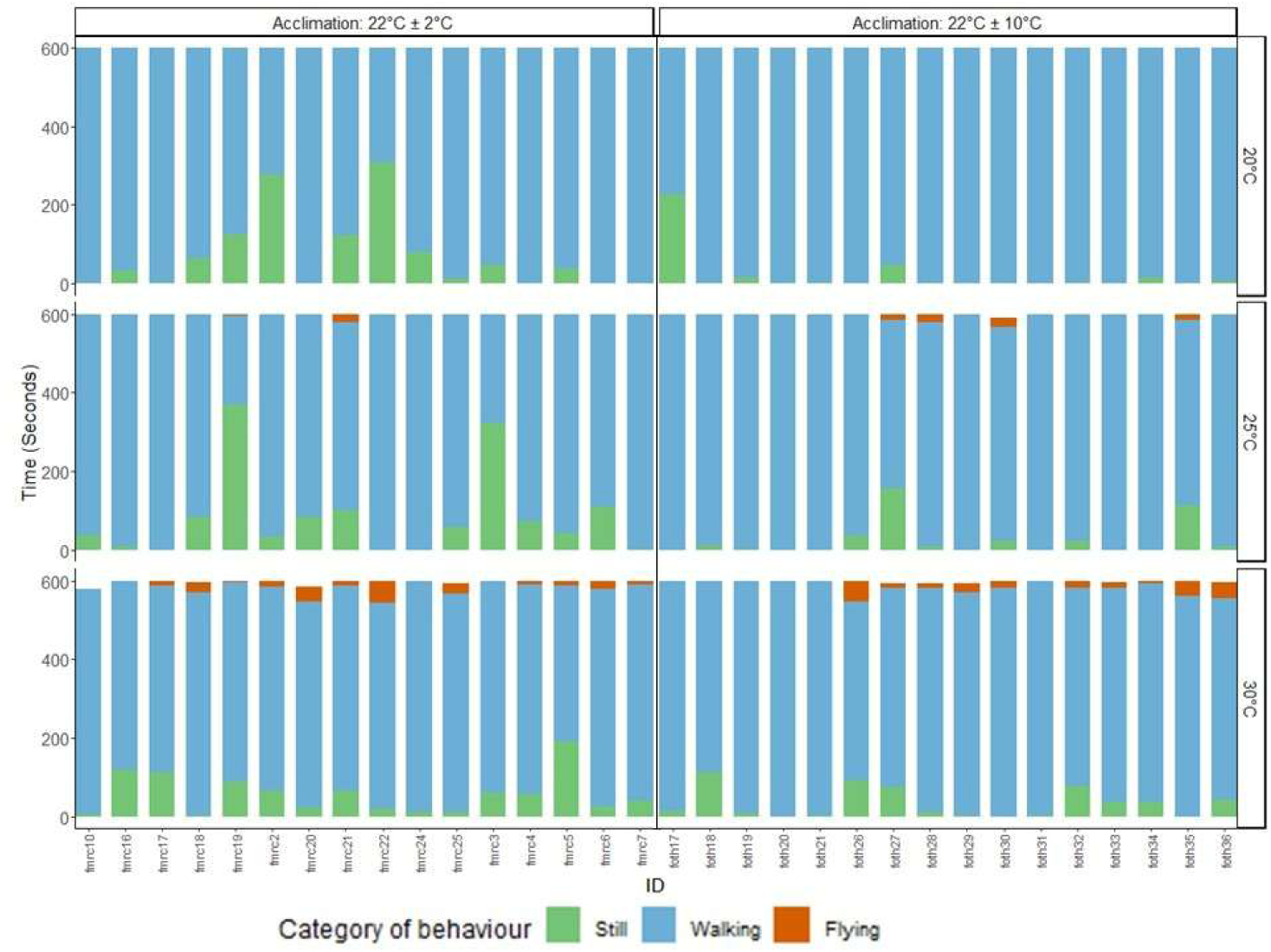
Activity during respirometry of *Onthophagus taurus* females. The X axis represents individuals and is grouped by fluctuation treatment: 22°C ± 2°C (left) and 22°C ± 10°C (right). The Y-axis shows the time in seconds that was spent on each behaviour. Colours represent behaviours: Green for still; blue for walking; and orange for flight. Statistics presented in text.

### Respirometry: energetic cost of locomotion

**Figure 3:**
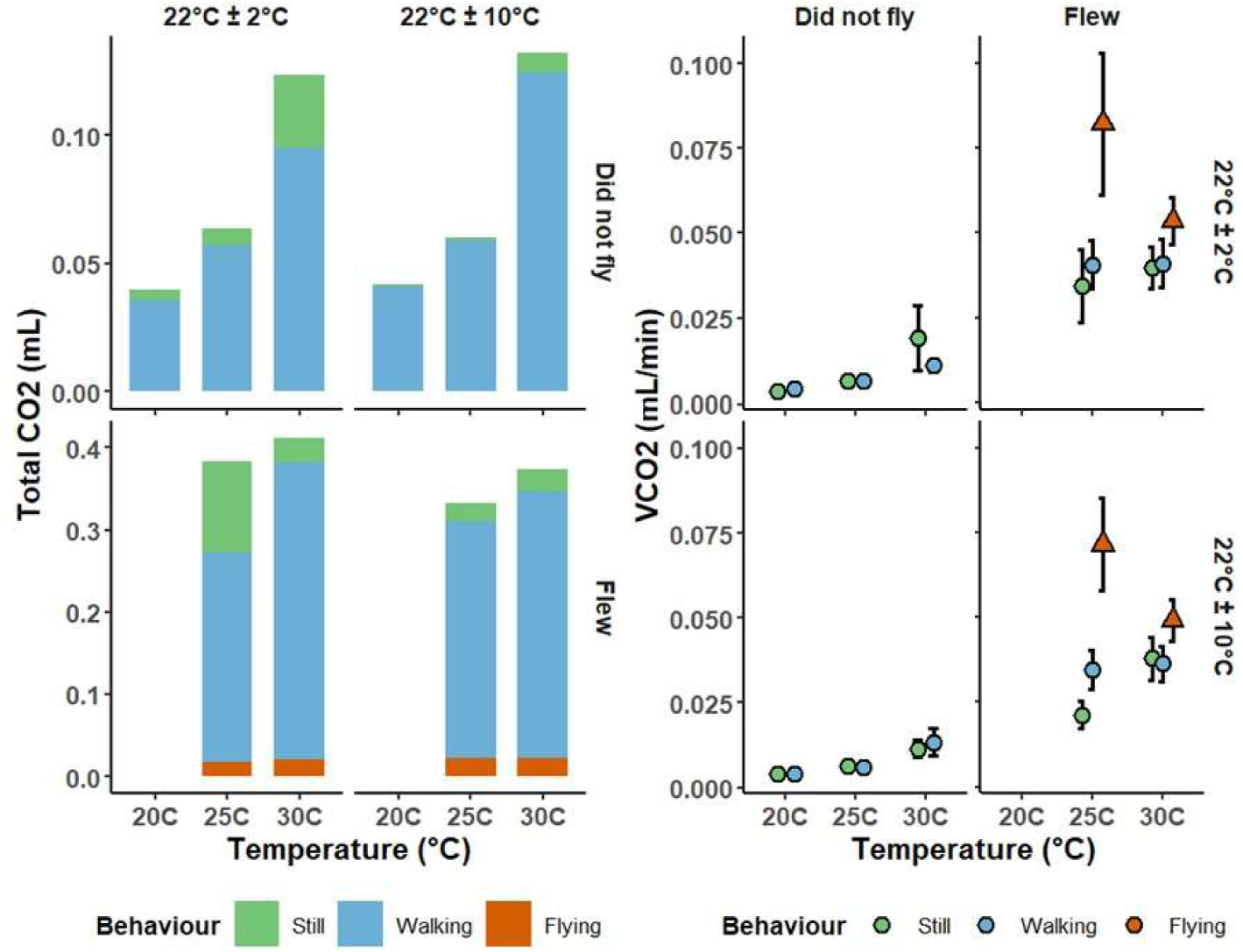
A) V̇CO _2_ during respirometry of female *Onthophagus taurus*. The X-axis represents trial temperature (i.e. air temperature in the chamber) (°C) and is grouped by fluctuation treatment: 22°C ± 2°C and 22°C ± 10°C. Colours represent the mean V̇CO _2_ per grouping. Beetles are grouped by fluctuation treatment and whether flight occurred at each trial temperature. B) Total CO _2_ in mL during respirometry of female *Onthophagus taurus*. The X-axis represents trial temperature (°C) and is grouped by fluctuation treatment: 22°C ± 2°C and 22°C ± 10°C. Colours represent the mean V̇CO_2_ per grouping. Beetles are grouped by fluctuation treatment and whether flight occurred at each trial temperature. Statistics presented in text.

Trial temperature (F_1,75.459_ = 23.524, *P* <0.005), mass (F_1,31.757_ = 18.740, *P* <0.005), and the amount of time spent flying (F_195.534_ = 375.725, *P* <0.005) were significant predictors of total CO_2_ production during the 10-minute measurement. While there was no effect of fluctuation treatment (F_1,43.014_ = 0.000, *P* = 0.998), the interaction between fluctuation treatment and the amount of time spent flying was a significant predictor of total CO_2_ production (F_1,89.543_ = 17.053, *P* <0.005).

No significant effect was found of fluctuation treatment on V̇CO_2_ for beetles in flight (F_1,24_ = 0.075, *P* = 0.787). Consistent with the literature, mass (F_1,24_ = 23.756, *P* <0.005) and trial temperature were significant predictors (F_1,24_ = 11.741, *P* < 0.005) of VCO2, however – as noted above – only six beetles flew below 30 °C. The time spent flying was a significant predictor of V̇CO_2_ while in flight, with greater energy expended per second in flight when the total flight time is lower (F_1,24_ = 14.222, *P* <0.005). For walking, the full model was dropped in favour of the reduced model omitting fluctuation treatment. Trial temperature (χ^2^ _(1)_ =198.280, *P* < 0.005) and mass (χ^2^ _(1)_ =6.851, *P* = 0.009) were significant predictors of V̇CO_2_. Similarly, the full model was dropped in favour of a reduced GLM excluding fluctuation treatment to test the effect of acclimation to thermal variability on V̇CO_2_ while beetles remained still. For beetles remaining still, trial temperature explained the majority of the variance (χ^2^ _(1)_ =156.915, *P* < 0.005), and mass was also a significant predictor (χ^2^ _(1)_ =5.767, *P* = 0.016). In summary fluctuation treatment does not explain differences in metabolic rate for any behaviour in our study.

### Mechanistic model

**Figure 5:**
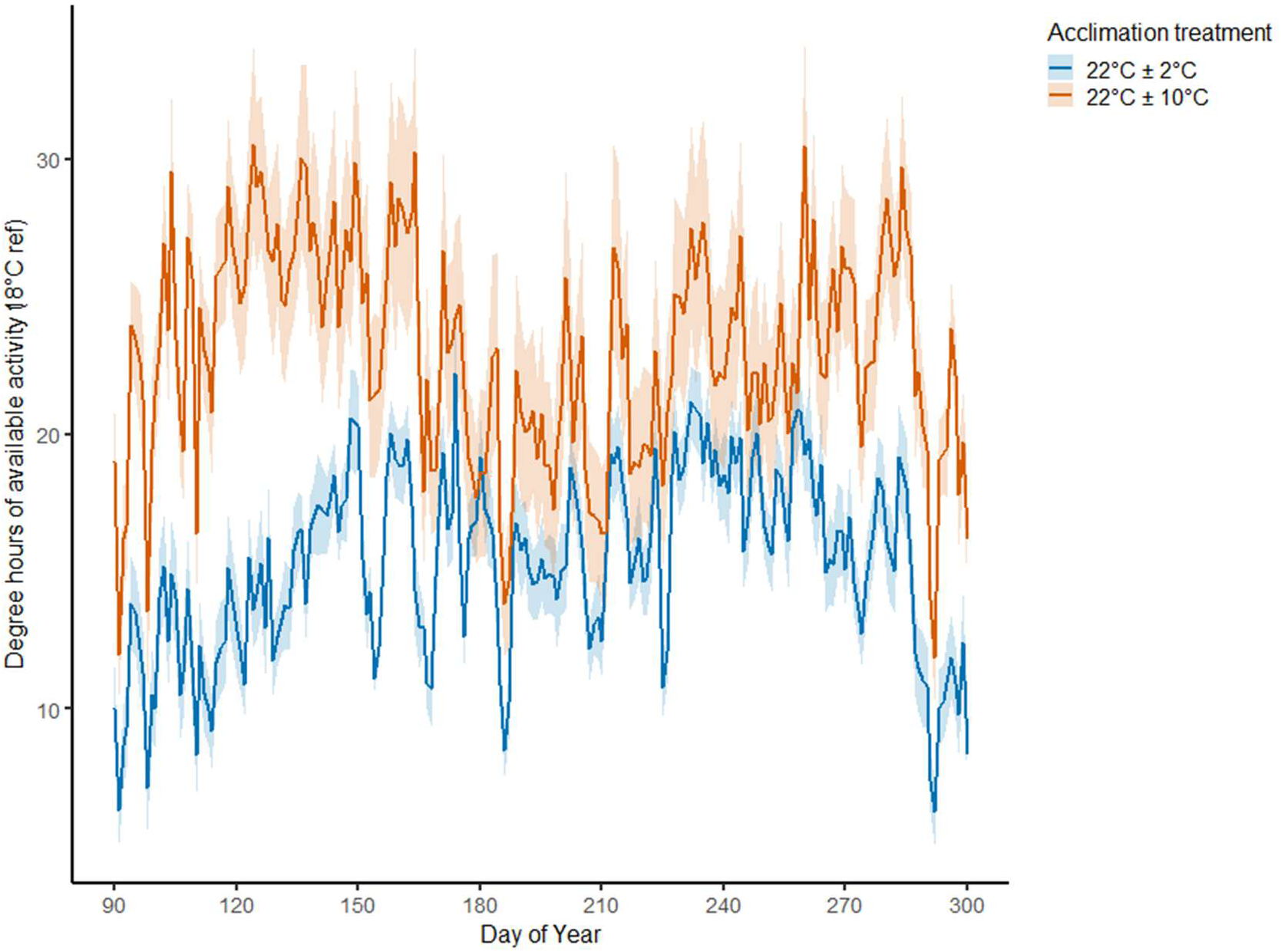
Predicted mean of total daily degree hours above 18 °C across individuals and years: 2018-2023, at Knoxville airport Tennessee US (35.81801°, -83.98573°), between day 90 and 300 of the year. Colours: high fluctuation (orange) and low fluctuation (blue) treatments.

**Figure 6:**
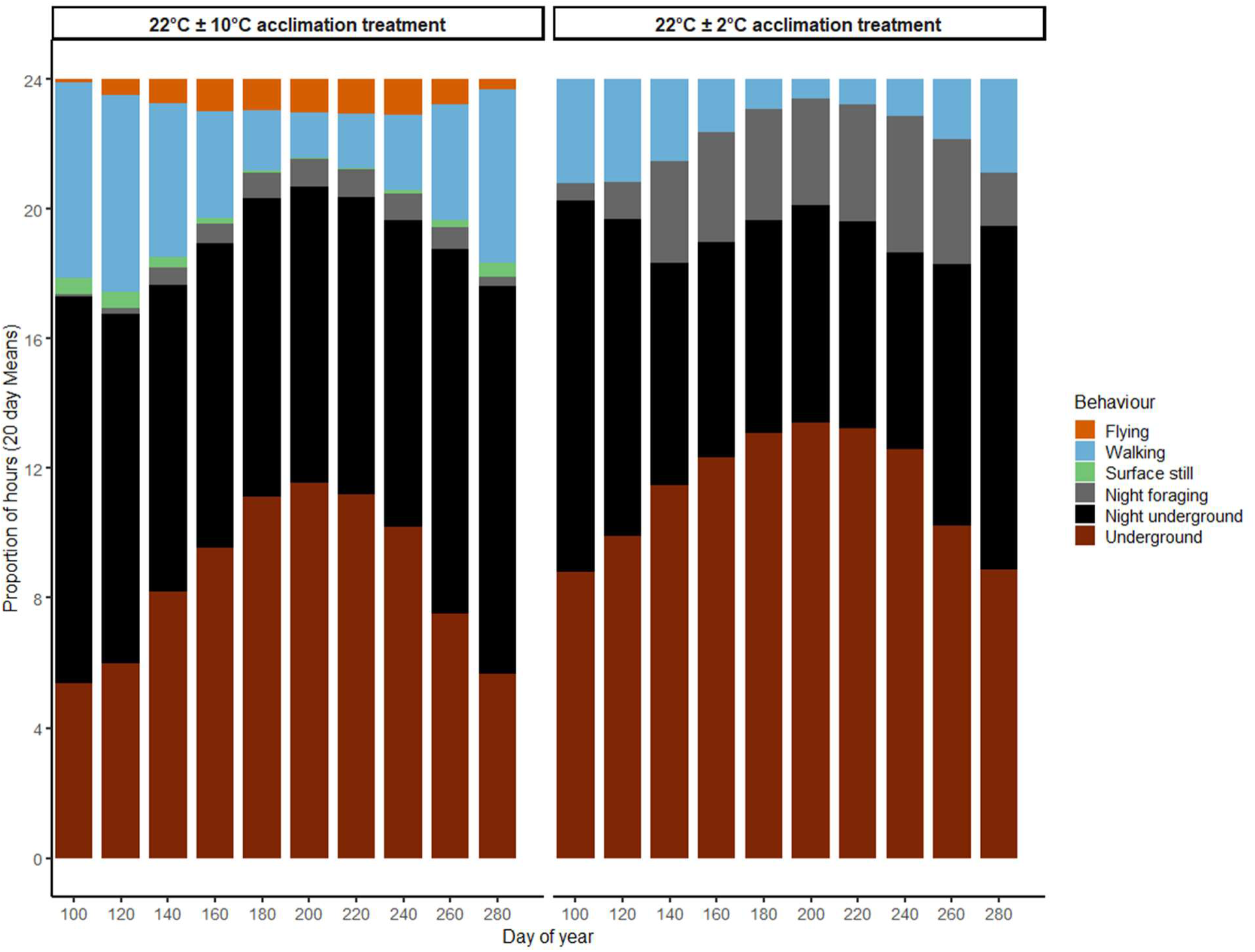
Predicted proportion of 24 hours with the opportunity for each behaviour averaged across individuals for the years: 2018-2023, at Knoxville Airport Tennessee US (35.81801°, -83.98573°), between day 90 and 300 of the year. Orange: flying, Blue: walking, Light green: Surface still, Gray: night foraging, Black: sheltering at night, Brown: daytime retreat. Statistics presented in text.

After examining modeled accumulated daily degree hours above the minimum foraging temperature across individuals and between acclimation treatments, we found no differences in total daily degree hours based on fluctuation treatment (χ^2^ _(1)_ =0.156, *P* = 0.693). However, both the day of year (χ^2^ _(2)_ =401.810, *P* < 0.005), and the interaction between day of year and fluctuation treatment were significant predictors of accumulated degree hours during activity (χ^2^ _(2)_ =475.562, *P* < 0.005), indicating seasonal patterns of accumulated degree hours while searching differed between fluctuation treatments. Notably, differences between fluctuation treatments were greater in the spring and autumn than in summer where high temperatures restricted diurnal search opportunities in both fluctuation treatments, leading to increased night searching and longer foraging hours in the low fluctuation group, and narrow activity windows at higher temperatures in those from the high fluctuation group. The narrower window of activity coincided with increased opportunities for flight that were not available to *O. taurus* from the low fluctuation group.

## Discussion

We provide evidence that acclimation to greater temperature fluctuations alters behaviour in *O. taurus* across a range of temperatures, thereby modifying energy expenditure during surface activity. Our results caution against relying only on individuals acclimated to constant temperatures when inferring energy expenditure from approaches that integrate behavioural and physiological data. As hypothesised, following acclimation to a high amplitude of thermal variability (22°C ± 10°C), *O. taurus* increases the intensity of activity over a narrower window of time compared to when maintained under a lower amplitude, at near-constant temperatures (22°C ± 2°C). Under acclimation to a high amplitude of thermal variability, *O. taurus* activity is restricted to rising diurnal temperatures, whereas nocturnal activity is seen in individuals acclimated to a low amplitude of variability. Acclimation to a high amplitude of variability increases overall locomotion and, in particular, flight activity at intermediate temperatures, which affects differences in energy expenditure. Flight was predicted only in individuals acclimated to high amplitudes of thermal variability and coincided with reduced time available for surface searching behaviour.

The prediction that, under increased amplitudes of thermal variability, daily activity is concentrated into a small period of time and time spent above ground is reduced, was supported. Beetles acclimated to greater thermal variability restricted surface activity to a narrow thermal window – similar to this species’ optimum range of temperatures for development (Floate et al., 2017). Activity was limited to the first period during the daily cycle that these temperatures were reached and only occurred during daylight, aside from in one individual. Comparatively, beetles acclimated to lower thermal variability displayed no consistent temporal pattern in their activity and were frequently active at night. Temperature and light are interacting regulators of circadian locomotory activity in many insect species (Beer & Helfrich-Förster, 2020). For instance, the nocturnal desert dwelling domino beetle (*Thermophilum duodecimguttatum*) differed in activity between induced thermal stability and variability under controlled conditions. Under thermally variable regimes activity would occur during the day to maintain activity at high temperatures (Constantinou & Cloudsley-Thompson 1985).

Plasticity in diel activity can accommodate differences in seasonal temperatures and microclimate availability. Diel activity has been shown to vary among habitats and seasons in the metallic brown harp ground beetle (*Poecilus chalcites*) with reduced diurnal activity in fields characterised by high variability compared with forest edge characterised by low variability (Willard & McCravy 2006).

Changes in diel activity can assist in reaching the optimum searching temperature or avoiding undesirable temperatures. In *O. taurus,* under greater thermal variability, burst activity occurred as temperatures were rising but stopped well before the maximum temperature was reached, suggesting that these dung beetles restrict activity to intermediate temperatures reducing searching opportunities. Circadian rhythm impacts activity when synchronised with higher amplitudes of thermal cycles.

The prediction that exposure to greater temperature fluctuations increases locomotive activity was supported. In both fluctuation treatments, flight events were more common at 30 °C than at 25 °C; this is consistent with the expectation that flight probability increases with temperature. This phenomenon is well documented in other Coleoptera; for example, in the lupine beetle *Sitona gressorius,* flight probability and speed were greatly affected by temperature, with higher temperatures associated with consistent activity (Hannigan et al., 2023). Metabolic rate during flight did not account for differences in energy expenditure during the trial. Metabolic rate during flight was higher at 25 °C than 30 °C. This could be explained by the reduced number of flight events within individuals that flew. Short and infrequent bursts of flight have a proportionately higher associated metabolic rate as the energy expended during flight is greatest during take-off (Oertili, 1989).

Contrary to what was predicted, we found no evidence of suppression of metabolic rate for active beetles, with differences in flight activity between fluctuation treatments explaining differences in energy expenditure. Previous studies of this species under similar amplitudes of variability found reduced metabolic rate in inactive beetles (Fleming et al., 2021). However, this was not the case here, likely because the previous studies on *O. taurus* held the animal in the dark and used a small respirometry chamber to minimise disturbances and measure resting metabolic rate. Our study used a large (500 mL) respirometry chamber floating in a water bath and an iridescent bulb to encourage activity and flight. Energy expended during walking and flight activity did not differ based on fluctuation treatment. However, there were insufficient occurrences of beetles remaining still at high temperatures following acclimation to the high fluctuation treatment. This prevented any meaningful comparison of resting metabolic rate at such temperatures, and therefore differences in findings are likely not conflicting.

Overall, differences in the energy expenditure of active *O. taurus* after acclimation to thermal variability was driven by flight events and body mass. This can be explained by how at higher temperatures, some of the time spent in stillness or walking encompasses energetic costs incurred as a consequence of pre-flight warm up, which were not accounted for separately in this study.

Previous work in other beetles has shown that the regulation of thoracic temperature independent of air temperature is greater in larger species but not explained by inter-individual variation during pre-flight warm up and post flight (Merrick & Smith 2004). As a result of pre-flight warm up costs, an increased number of flights separated over time and longer periods of time spent in preparation (regardless of if flight eventually occurs), incurs greater total energetic costs.

The model shows that under environmental temperatures from where the species were collected, acclimation to thermal variability restricts summer activity time and accumulated daily degree hours. Activity concentrated at higher temperatures likely increases search speed including an allowance of opportunities for flight. Differences in degree hours are more apparent in spring and autumn than summer, although differences in behaviour are greater during summer. To our knowledge, this is the first study that focuses on differences in locomotive choices in a facultative flyer as a consequence of acclimation to increased thermal variability. Gillespie *et al*, (2017) observed flight activity period and peak flight activity across 588 beetle species in Norway over a 13 year period. They found smaller species were more abundant and flew for longer periods in the summer with flight activity peaking in midsummer. Smaller species, such as *Onthophagus taurus*, had longer temporal windows than larger species, and flight length decreased in cooler years.

Comparatively, flight length increased in cooler years for larger species. A high amount of midsummer activity fits with the species breeding season. Previous work has shown that in *O. taurus*, brood ball production is associated with mean temperature, and while possible in spring, is largely confined to summer (Fleming et al., 2024). In our study, differences in degree hours were lowest during the summer, suggesting reduced opportunity for searching behaviour.

To conclude, in a temperate dung beetle, exposure to greater thermal variability shapes diel patterns in activity and increases locomotive intensity when active. Our study highlights the need for laboratory studies to include fluctuation treatments that reflect the range of temperatures experienced in the field. When parameterising a mechanistic model using the results of this study, inferences reliant on information collected from insects held at a constant temperature would have been misleading and underestimated the capacity of individuals to fly in the field.

## Notes

https://osf.io/8km7x/overview

### Competing Interest Statement

The authors have declared no competing interest.

### Funding

- U.S. National Science Foundation

