## Supplemental tables and figures for "Acclimation to thermal variability increases the intensity of activity and alters the activity window in the temperate dung beetle *Onthophagus taurus*"

**Supplemental information**

Table S1: Incubator temperature and light and dark cycles for *O. taurus* beetles collected in Knoxville Tennessee, USA and acclimated to either 22°C ± 2°C or 22°C ± 10°C.

| **Time** | **Temperature: 22°C ± 2°C** | **Temperature 22°C ± 10°C** | **LD cycle** |
|---|---|---|---|
| 23:00 | 21.3 | 22 | Dark |
| 01:00 | 20.7 | 18.7 | Dark |
| 03:00 | 20 | 15.3 | Dark |
| 05:00 | 20.7 | 12 | Dark |
| 07:00 | 21.3 | 15.3 | Light |
| 09:00 | 22 | 18.7 | Light |
| 11:00 | 22.7 | 22 | Light |
| 13:00 | 23.3 | 25.3 | Light |
| 15:00 | 24 | 28.7 | Light |
| 17:00 | 23.3 | 32 | Light |
| 19:00 | 22.7 | 28.7 | Light |
| 21:00 | 22 | 25.3 | Light |

Table S2:

Anova results for the model predicting the time spent still above ground of *O. taurus* beetles collected in Knoxville Tennessee, USA and acclimated to either 22°C ± 2°C or 22°C ± 10°C. ΔAIC of 1.31, giving support for the limited model relative to the full model.

| | Chisq | Df | Pr(>Chisq) |
|---|---|---|---|
| (Intercept) | 6.5044 | 1 | 0.01076 |
| Incubator temperature | 5.9007 | 1 | 0.01514 |
| LD cycles | 3.7561 | 1 | 0.05261 |
| Incubator temperature × LD cycles | 4.0343 | 1 | 0.04458 |

Table S3:

Anova results for the model predicting the time spent moving above ground of *O. taurus* beetles collected in Knoxville Tennessee, USA and acclimated to either 22°C ± 2°C or 22°C ± 10°C. ΔAIC of full model relative to the next best model = 1.78.

| | Chisq | Df | Pr(>Chisq) |
|---|---|---|---|
| (Intercept) | 1.7142 | 1 | 0.190447 |
| Incubator temperature | 14.1473 | 1 | 0.000169 |
| LD cycles | 5.9398 | 1 | 0.014802 |
| Incubator temperature × LD cycles | 6.3230 | 1 | 0.011918 |
| Fluctuation treatment | 3.8887 | 1 | 0.048611 |

Table S4: Anova results for the GLM predicting time spent in flight during the respirometry measurements of *O. taurus* beetles collected in Knoxville Tennessee, USA and acclimated to either 22°C ± 2°C or 22°C ± 10°C. ΔAIC of full model relative to the next best model = 1.86.

| | Chisq | Df | Pr(>Chisq) |
|---|---|---|---|
| (Intercept) | 9.6446 | 1 | 0.0018991 |
| Chamber temperature | 14.8678 | 1 | 0.0001153 |
| Mass immediately before respirometry | 2.8470 | 1 | 0.0915449 |
| Fluctuation treatment | 6.0626 | 1 | 0.0138075 |
| Chamber temperature × Fluctuation treatment | 5.4784 | 1 | 0.0192530 |

Table S5: Anova results for the GLM predicting walking behaviour during the respirometry measurements of *O. taurus* beetles collected in Knoxville Tennessee, USA and acclimated to either 22°C ± 2°C or 22°C ± 10°C. ΔAIC of full model relative to limited model = 4.68.

| | Chisq | Df | Pr(>Chisq) |
|---|---|---|---|
| (Intercept) | 2447.8087 | 1 | < 2.2e-16 |
| Chamber temperature | 0.5171 | 1 | 0.472060 |
| Mass immediately before respirometry | 0.2247 | 1 | 0.635452 |
| Fluctuation treatment | 6.9262 | 1 | 0.008494 |

Table S5: Anova results for the GLM predicting still behaviour during the respirometry measurements of *O. taurus* beetles collected in Knoxville Tennessee, USA and acclimated to either 22°C ± 2°C or 22°C ± 10°C. ΔAIC of full model relative to limited model = 4.99.

| | Chisq | Df | Pr(>Chisq) |
|---|---|---|---|
| (Intercept) | 7.7757 | 1 | 0.005295 |

| | | | |
|---|---|---|---|
| Chamber temperature | 0.0656 | 1 | 0.797864 |
| Mass immediately before respirometry | 0.1100 | 1 | 0.740157 |
| Fluctuation treatment | 7.5321 | 1 | 0.006061 |

Table S6: Linear model results of predicting total $CO_2$ production during respirometry measurements for *O. taurus* beetles collected in Knoxville, Tennessee, USA and acclimated to 22°C ± 2°C and 22°C ± 10 °C. ΔAIC of full model was 14.86 relative to the next best reduced model.

| | Estimate | Standard error | df | t value | Pr(>\|t\|) |
|---|---|---|---|---|---|
| (Intercept) | -0.2990160 | 0.0619108 | 89.4900610 | -4.830 | 5.61e-06 |
| Mass immediately before respirometry | 1.5975345 | 0.3913590 | 28.6607354 | 4.082 | 0.000326 |
| Time spent in flight | 0.0148260 | 0.0009013 | 85.3028957 | 16.449 | < 2e-16 |
| Fluctuation treatment: 22°C ± 10 °C | 0.0002125 | 0.0161452 | 38.3759072 | 0.013 | 0.989567 |
| Chamber temperature | 0.0097105 | 0.0020482 | 72.2782562 | 4.741 | 1.04e-0 |
| Time spent in flight × Fluctuation treatment: 22°C ± 10 °C | -0.0045932 | 0.0011332 | 84.2989170 | -4.053 | 0.000112 |

Table S7: Anova results for the LM predicting $\dot{V}CO_2$ while in flight during the respirometry measurements of *O. taurus* beetles collected in Knoxville Tennessee, USA and acclimated to either 22°C ± 2°C or 22°C ± 10°C. ΔAIC of full model was 9.55 relative to the next best reduced model.

| | df | Sum sq | Mean Sq | F value | Pr(>F) |
|---|---|---|---|---|---|
| Chamber temperature | 1 | 0.0026371 | 0.0026371 | 11.7410 | 0.0022087 |
| Mass immediately before respirometry | 1 | 0.0055604 | 0.0055604 | 24.7561 | 4.423e-05 |
| Fluctuation treatment: 22°C ± 10 °C | 1 | 0.0000168 | 0.0000168 | 0.0749 | 0.7866681 |
| Time spent in flight | 1 | 0.0031943 | 0.0031943 | 14.2217 | 0.0009375 |

Table S8: Anova results for the GLM predicting $\dot{V}CO_2$ while walking during the respirometry measurements of *O. taurus* beetles collected in Knoxville Tennessee, USA and acclimated to either 22°C ± 2°C or 22°C ± 10°C. ΔAIC = 1.96 giving support for the limited model relative to the full model.

| | Chisq | Df | Pr(>Chisq) |
|---|---|---|---|
| Chamber temperature | 198.2800 | 1 | < 2.2e-16 |
| Mass immediately before respirometry | 6.8511 | 1 | 0.008859 |

Table S9: Anova results for the GLM predicting $\dot{V}CO_2$ while still during the respirometry measurements of *O. taurus* beetles collected in Knoxville Tennessee, USA and acclimated to either 22°C ± 2°C or 22°C ± 10°C. ΔAIC = 1.99 giving support for the limited model relative to the full model.

| | Chisq | Df | Pr(>Chisq) |
|---|---|---|---|
| Chamber temperature | 156.9145 | 1 | < 2e-16 |

| | | | |
|---|---|---|---|
| Mass immediately before respirometry | 5.7676 | 1 | 0.01632 |

*Details of Biophysical models*

*Microclimate Model*

Our models were built based on Knoxville Airport, Tennessee, US (35.81801°, -83.98573°).

Table S10:

| Variable | |
|---|---|
| Reflectance | 0.15 decimal % |
| Slope | 0° |
| Aspect | North facing |
| Node depth | 0, 2.5, 5, 10, 15, 20, 30, 50, 100, 200 CM |
| Shade range | 0 to 50 % |
| local height at which air temperature, wind speed and humidity are to be computed | 0.01 M |
| Run moist | 1 |
| Clear sky | Observed cloud cover included |
| Run gads | Global soil database R version |

| | |
|---|---|
| Soil grids | Soil hydraulic properties queried from soilgrids.org |

Table S11: Ectotherm model parameters taken from the literature or estimated that effected the heat budget of the organism

| **Component** | **Value** | **Species** | **Source** |
|---|---|---|---|
| Critical thermal minimum | 1.2°C | *Acrossus depressus* | Birkett *et al.,* 2018 |
| Orientation to the sun (posture) | Orient perpendicular to the sun | *Onthophagus taurus* | Dacke *et al*., 2014 |
| Colour morph | 85% - 95% absorptivity | *Geotrupes auratus* | Watanabe *et al.,* 2002 |
| Shade seeking | Yes | *Scarabaeus (Kheper) lamarcki* | Smolka *et al.,* 2012 |
| Burrow | Yes | *Onthophagus taurus* | Mamantov & Sheldon., 2021 |
| Climb | No, simulating flat landscape | | |
| Mindepth | $2^{nd}$ Node | *Onthophagus taurus* | Mamantov & Sheldon., 2021 |
| Max depth | $6^{th}$ Node | *Onthophagus taurus* | Mamantov & Sheldon., 2021 |

| | | | |
|---|---|---|---|
| Activity allowed in rain | Yes | Multiple assemblages of dung beetle | Sun *et al.,* 2003 |
| Transient | No | | |
| Surface area of eyes | 1% | | Estimation |
| Surface area of mouth | 2% | | Estimation |
| Temperature difference between inhaled and exhaled air | 0.1 (°C) | | Estimation |
| Warming is a signal for emergence | Yes | | Assumption |
| Panting as a multiplier of breathing rate | Off | | Ectotherm |
| Percentage of body in contact with surface | 1% | | Estimation |
| Percentage shade increment | 1% | | |
| Live | Full behaviour | | |

*Ectotherm Model*

Within the NicheMapR models, *O. taurus* mass is the organism-specific variable which shifts between treatments and controls for changes in body temperature. We took other parameters from previous research when available (Table S11). Where information for *O. taurus* is unavailable, we used data from most morphologically similar species of dung beetle. The model calculated behavioural thermoregulation in three categories: inactivity, basking, and foraging. Inactive hours are periods where *O. taurus* burrow to avoid surface conditions.

*Model parameters taken from the literature*

Some variables were assumed to be negligible due to the size of the organism and accordingly given very low values, these included: surface area taken up by open eyes and mouth, temperature difference for expired and inspired air, and percentage of the animal's surface in contact with the substrate. *Onthophagus taurus* is ellipsoid shaped, and orients perpendicular to the sun (Pokhrel *et al*., 2020 & Dacke *et al*., 2014).

*Onthophagus taurus* exhibits significant variations in its body colour (Parzer *et al.,* 2018). Solar absorptivity for the species had to be estimated using the available species of dung beetle with the most similar morphology (*Geotrupes auratus*; Watanabe *et al.,* 2002). While the species have different colour morphs, the difference in body temperature resulting from incremental changes to the reflectance value were negligible. This is a steady state model, and we made the assumption that beetles will select the warmer conditions whether that be on the surface or not within the temperature range at which activity occurred during the experimental trials.

The beetle will need the capacity to do so and accordingly we have estimated the temperature at which the species will experience cold stupor. We could find no examples in the literature of dung beetles expressing behaviour contrary to that and the selected temperature is in line with previous work on lizards using NicheMapR (Kearney *et al* 2018). Chill coma varies from 3.1 to 1.6 °C above the critical thermal minimum for the species *Sulcophanaeus imperator* and *Sulcophanaeus batesi* respectively (Verdu *et al.,* 2019). We have used the former species, that reaches chill coma at a higher temperature, to compensate for *O. taurus* becoming increasingly lethargic at lower temperatures.

*Model limitations*

Our model predicts the heat budget of the organism when searching behaviour occurs, but we do not consider energetic cost directly as amount of time spent in flight, length of flight, and length of time spent searching is beyond the scope of our study. Instead, we present when these behaviours can occur based on the behavioural data collected for the model. Our assumptions are based on behaviour recorded from measurements taken in a lab setting and further field studies are needed.

*Biophysical model statistics*

Table S12: Anova results for the GLM predicting degree hours of foraging for *O. taurus* beetles collected in Knoxville Tennessee, USA and acclimated to either 22°C ± 2°C or 22°C ± 10°C. The full model had an ΔAIC=472 relative to the next best reduced model.

| | Chisq | Df | Pr(>Chisq) |
|---|---|---|---|
| (Intercept) | 75.5919 | 1 | <2e-16 |
| Daily degree hours spent searching | 401.8104 | 1 | <2e-16 |
| Fluctuation treatment | 0.1562 | 1 | 0.6926 |

| | | | |
|---|---|---|---|
| Daily degree hours spent searching × Fluctuation treatment | 475.5619 | 1 | <2e-16 |

*Citations for supplementary materials*


Birkett, A. J., Blackburn, G. A., & Menéndez, R. (2018). Linking species thermal tolerance to elevational range shifts in upland dung beetles. *Ecography, 41*(9), 1510–1519. https://doi.org/10.1111/ecog.03399

Dacke, M., el Jundi, B., Smolka, J., Byrne, M., & Baird, E. (2014). The role of the sun in the celestial compass of dung beetles. *Philosophical Transactions of the Royal Society B: Biological Sciences, 369*(1636), 20130036. https://doi.org/10.1098/rstb.2013.0036

Kearney, M. R., Munns, S. L., Moore, D., Malishev, M., & Bull, C. M. (2018). Field tests of a general ectotherm niche model show how water can limit lizard activity and distribution. *Ecological Monographs, 88*(4), 672–693. https://doi.org/10.1002/ecm.1326

Mamantov, M. A., & Sheldon, K. S. (2021). Behavioural responses to warming differentially impact survival in introduced and native dung beetles. *Journal of Animal Ecology, 90*(1), 273–281. https://doi.org/10.1111/1365-2656.13363

Parzer, H. F., Polly, P. D., & Moczek, A. P. (2018). The evolution of relative trait size and shape: Insights from the genitalia of dung beetles. *Development Genes and Evolution, 228*(2), 83–93. https://doi.org/10.1007/s00427-018-0603-2

Pokhrel, M. R., Cairns, S. C., & Andrew, N. R. (2020). Dung beetle species introductions: When an ecosystem service provider transforms into an invasive species. *PeerJ, 8*, e9872. https://doi.org/10.7717/peerj.9872

Smolka, J., Baird, E., Byrne, M. J., el Jundi, B., Warrant, E. J., & Dacke, M. (2012). Dung beetles use their dung ball as a mobile thermal refuge. *Current Biology, 22*(20), R863–R864. https://doi.org/10.1016/j.cub.2012.08.043

Sun, W., Tang, W., Wu, Y., He, S., & Wu, X. (2023). The influences of rainfall intensity and timing on the assemblage of dung beetles and the rate of dung removal in an alpine meadow. *Biology, 12*(12), 1496. https://doi.org/10.3390/biology12121496

Verdú, J. R., Cortez, V., Oliva, D., & Giménez-Gómez, V. (2019). Thermoregulatory syndromes of two sympatric dung beetles with low energy costs. *Journal of Insect Physiology, 118*, 103945. https://doi.org/10.1016/j.jinsphys.2019.103945

Watanabe, T., Tanigaki, T., Nishi, H., Ushimaru, A., & Takeuchi, T. (2002). A quantitative analysis of geographic color variation in two *Geotrupes* dung beetles. *Zoological Science, 19*(3), 351–358. https://doi.org/10.2108/zsj.19.351
